# Electroporation of short hairpin RNAs for rapid and efficient gene knockdown in the starlet sea anemone, *Nematostella vectensis*

**DOI:** 10.1101/422113

**Authors:** Ahmet Karabulut, Shuonan He, Cheng-Yi Chen, Sean A. McKinney, Matthew C. Gibson

**Affiliations:** Stowers Institute for Medical Research, Kansas City, Missouri, USA

## Abstract

A mechanistic understanding of evolutionary developmental biology requires the development of novel techniques for the manipulation of gene function in phylogenetically diverse organismal systems. Recently, gene-specific knockdown by microinjection of short hairpin RNA (shRNA) has been applied in the sea anemone *Nematostella vectensis,* a cnidarian model organism. Due to the unusual architecture of the cnidarian microRNA processing pathway, the shRNA approach is unusually effective for sequence-specific knockdown of a gene of interest. However, the time- and labor-intensive process of microinjection limits access to this technique and its application in large scale experiments. To address this issue, here we present an electroporation protocol for shRNA delivery into *Nematostella* eggs. This method leverages the speed and simplicity of electroporation, enabling users to manipulate gene expression in hundreds of *Nematostella* eggs or embryos within minutes. We provide a detailed description of the experimental procedure, including reagents, electroporation conditions, preparation of *Nematostella vectensis* eggs, and follow-up care of experimental animals. Finally, we demonstrate the knockdown of several endogenous and exogenous genes with known phenotypes and discuss the potential applications of this method.

## INTRODUCTION

The starlet sea anemone, *Nematostella vectensis,* is an established cnidarian model organism for *in vivo* studies of developmental processes^1,2^. *Nematostella* has several unique advantages to study evolutionarily conserved pathways that regulate gene function in metazoans^3^. First, *Nematostella* is a broadcast spawner, allowing massive numbers of gametes to be collected in a single spawn. Second, *Nematostella* features relatively rapid embryonic and larval development, and the embryos are easy to visualize and manipulate. Third, the *Nematostella* genome has been sequenced, with transcriptomic profiles available for a variety of developmental stages and tissues^4^. Last but not least, comparative genomic analysis indicates that despite a simple adult body plan, a large number of bilaterian genes and signaling pathways are conserved in *Nematostella*^3^. For example, *Nematostella* possesses the majority of known bilaterian signaling cascades including the Wnt^5^, Notch^6,7^, Bone Morphogenetic Protein (BMP) ^8^, Fibroblast Growth Factor (FGF)^9,10^, and Hedgehog pathways^11^. Furthermore, a high percentage of introns and the overall intron-exon structure of human genes are also relatively conserved in *Nematostella*^12,13^. For these and additional reasons, *Nematostella* is an outstanding model organism for comparative studies of cellular, developmental and evolutionary biology.

Since the early 2000s, *Nematostella* researchers have developed several protocols including maintenance and spawning procedures^14,15^, isolation of protein and nucleic acids^16^, *in situ* hybridization^17,18^, microinjection and morpholino-based gene knockdown^19^, meganuclease-mediated transgenesis ^20^ and CRISPR/Cas9-based gene editing ^21^. Recently, we developed a gene knockdown method based on microinjection of gene-specific shRNA into *Nematostella* eggs (He et al., *in press* 2018). This method allows robust and cost-effective gene silencing, but the reliance on microinjection introduces experimental limitations, particularly the number of eggs that can be injected and later recovered. In addition, the precision control of injection volume is difficult. Thus, a more efficient method for shRNA delivery would be ideal for medium-and high-throughput applications. As an alternative to microinjection, electroporation has been utilized for biomacromolecule delivery into *Hydra* ^*22*^ and *Ciona*^*23,24*^. Here, we report an optimized method that combines the advantages of electroporation with the specificity of the shRNA-based gene knockdown approach in *Nematostella*. Unlike microinjection, this electroporation protocol can be performed by novice users, requires minimal effort, and can quickly deliver shRNA to a large number of eggs simultaneously.

## EXPERIMENTAL DESIGN

### Synthesis and preparation of shRNA

The shRNA-mediated knockdown method is based on *in vitro* transcription (IVT) of shRNA from a template DNA using T7 RNA Polymerase. An illustration of the IVT reaction is provided in **(Fig 1a).** We used a 19bp gene targeting motif size, which has been found to be optimal and cost-effective for gene knockdown^25^. DNA templates for IVT were assembled by direct annealing of complimentary forward and reverse oligos. Primers were mixed 1:1 to a final concentration of 50μM per primer and denatured at 98°C for 5 minutes. Upon annealing at 24°C, the assembled duplex was kept at room temperature for immediate use as the template for IVT. We typically allowed transcription reactions to continue for 5 to 7 hours. DNase was then added to remove the template, and the reaction mix was purified using the Ampliscribe™ T7-Flash™ Transcription Kit (Invivogen, Inc). A single IVT reaction using this kit typically yielded 2-3 μg/μl purified shRNA in a 30μl elution volume as assessed by a spectrophotometer (we considered purity adequate at a 260/280 ratio above 1.80). In general, we have observed that a lower purity is not toxic to eggs when used for electroporation. Considering the typical yield, a researcher could potentially bypass the purification step and use the IVT product directly for electroporation if the aim is a large phenotypic screen. The shRNA templates used in this study can be seen in **Table 1**.

**Table 1:**
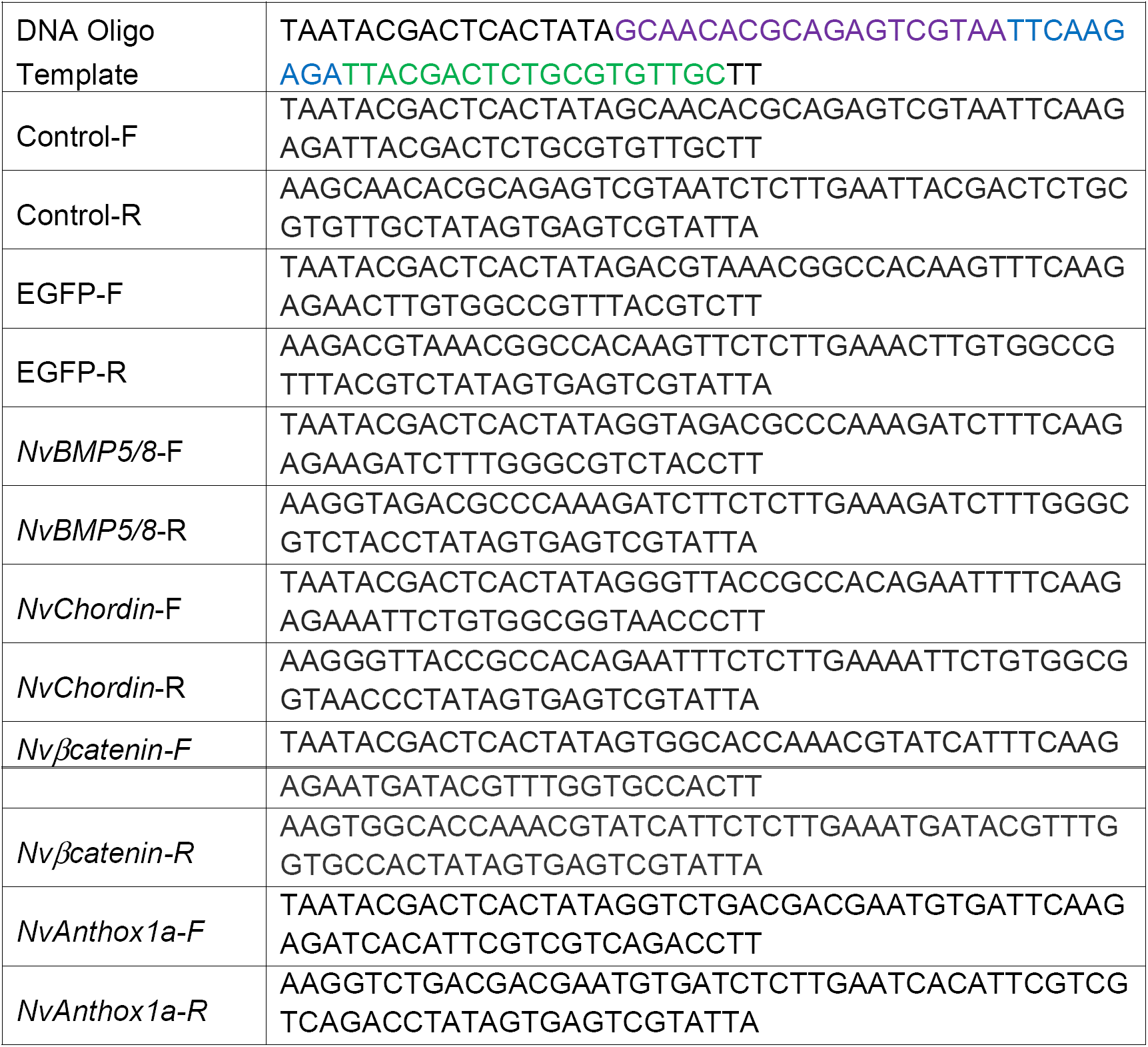
Template sequences used for shRNA synthesis. The DNA Oligo Template-F sequence indicates the positions of the 5’ T7 promoter (**black**), 19 nucleotide motif (magenta), a linker loop (blue), corresponding anti-sense motif (green) and two thymidine bases at 3’.

**Figure 1.**
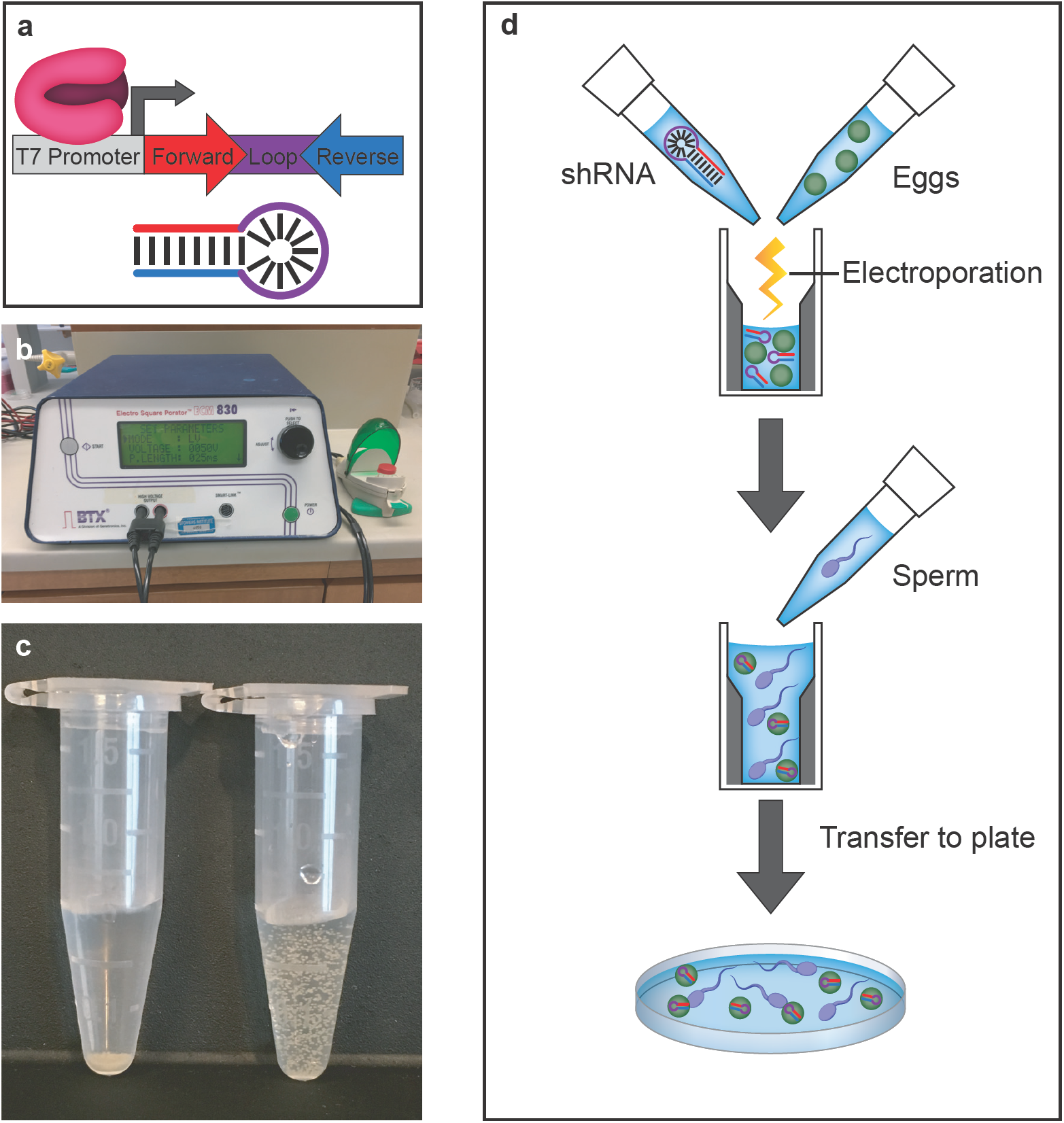
The procedure for electroporation of shRNA into *Nematostella* eggs. **(a)** *In vitro* transcription using T7 RNA Polymerase and the template. (**b**) EMM 830 square wave electroporator (BTX instruments) with attached Gene Pulser Xcell ShockPod Cuvette Chamber (Bio-rad) and 4mm Cuvette (Mirus). LCD display indicates Electroporation settings (50 Volts, 25ms (single pulse). (**c**) Suspension of eggs using 15% Ficoll PM 400 in 12 p.p.t salt water. Left: Eggs in 12 p.p.t salt water. Right: Eggs suspended in 12 p.p.t. salt water with 15% Ficoll PM400. **(d)** Illustration of the electroporation procedure.

### Electroporation of shRNA

*Nematostella* eggs have a higher density than sea water and tend to settle at the bottom of an electroporation cuvette, potentially resulting in non-uniform electroporation. To solve this problem, we suspended eggs in a 15% Ficoll PM 400 solution before transfer to the electroporation cuvette (**Fig. 1b**). shRNA was then added to the cuvette and mixed by gentle shaking for 10 seconds. Following these simple steps, eggs should be evenly suspended in the cuvette.

Once eggs are distributed in the cuvette, electroporation uses pulses of electricity to temporarily form pores throughout the plasma membrane. During this temporal window, shRNA can be efficiently delivered into the egg. To exploit this, we used a standard laboratory electroporation setup to knock down genes of interest using shRNA **(Fig. 1a).** Electroporation efficiency depends on the parameters of voltage, pulse duration, and number of pulses. We therefore performed multivariate experiments to determine electroporation efficiency by evaluating embryo survival and subsequent knockdown penetrance for the conserved Wnt pathway component *Nvβ-Catenin* (**Fig. 1a, Supplemental Table 1**). Optimal parameters were determined to be 50 volts delivered in a single 25 millisecond pulse in 4mm cuvettes (Mirus Bio) using an Electroporator ECM830 (Genentronics Inc.; **Fig. 1a**). These settings provided efficient shRNA delivery and gene knockdown with an acceptable amount of egg loss (15-20%). The overall workflow is illustrated in (**Fig. 1d)**.

Factors that might affect egg recovery include the number of unfertilized eggs per cuvette, the age of the eggs (hours post-spawn), and the care given to post-electroporation handling. In our experience, eggs from a fresh spawn are necessary for optimal survival. After electroporation, eggs tend to display abnormal morphologies and frequently exhibit a single protrusion on one side. These typically recover but are fragile and should be immediately transferred to a petri dish, fertilized, and incubated at low density without disruption. At higher densities oocytes may fuse inside the cuvette and hence the number of eggs (ideally around 300-600 per cuvette) should be monitored. We also found that electroporation can be performed with fertilized eggs, which do not fuse in the cuvette.

## VALIDATION AND RESULTS

Following electroporation, the extent of target gene knockdown hypothetically depends on the shRNA dose. To test this, we electroporated either *EGFP shRNA* or an shRNA scramble control into *Actin-EGFP* transgenic animals and analyzed EGFP protein levels by fluorescent microscopy (**Fig. 2a-i**). In this case, the optimal EGFP shRNA concentration was approximately 300ng/μl as higher concentrations did not produce a stronger knockdown effect (**Fig 2f**). However, based on our experience with multiple experiments, the optimal shRNA concentration is gene-dependent. In general, *Nematostella* eggs tolerate high concentrations of shRNA. Here, we electroporated up to 500ng/μl shRNA and did not observe any obvious toxicity (**Fig. 2j-r**). In other experiments, concentrations as high as 900ng/μl were used without any obvious effects on embryo survival **(e.g. Figure 6a, c).**

**Figure 2.**
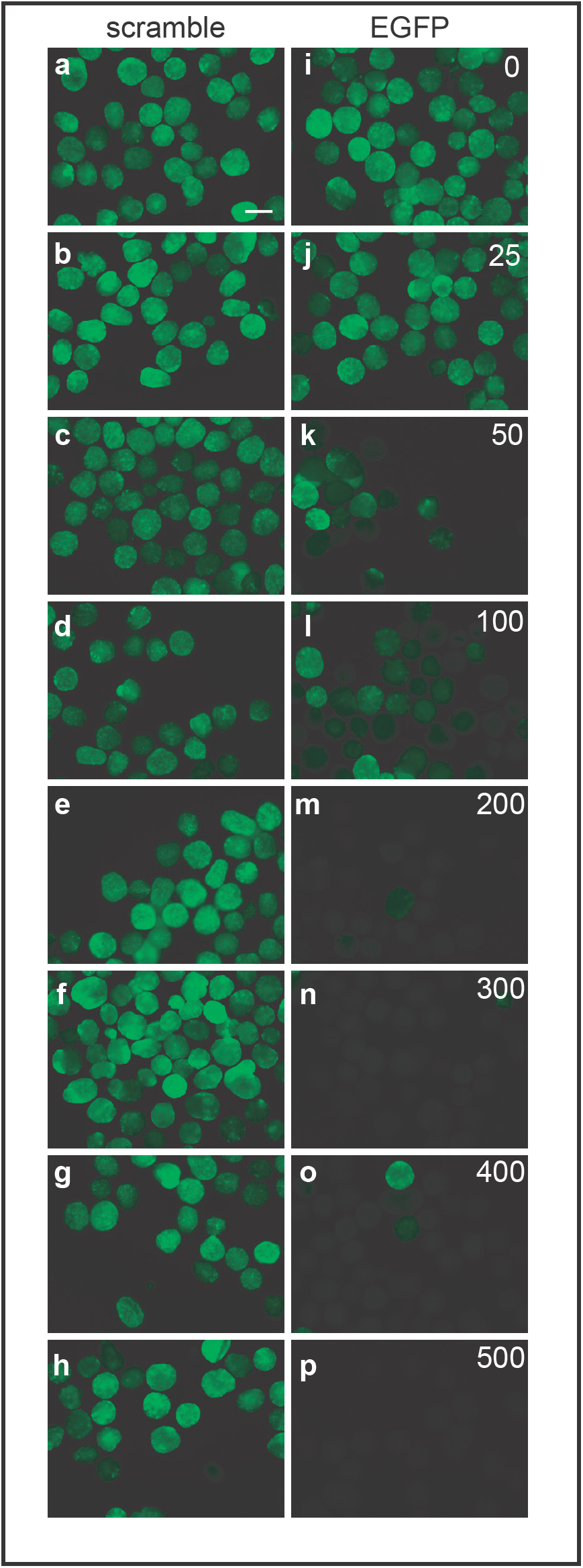
Typical dose response results of varying concentrations of either EGFP shRNA or scramble (control) shRNA after electroporation into *Actin-EGFP* transgenic animals. Images were acquired 72 hours after electroporation.(**a-h**) Incremental doses of scramble (control) shRNA (ng/μl). (**i-p**) Corresponding incremental doses of EGFP shRNA. (Scale Bar = 200 μm)

To quantify the level of gene-specific knockdown at varying doses of shRNA, we next electroporated 100, 200 and 400ng/μl *EGFP shRNA* into *Actin-EGFP* transgenic animals. EGFP signal was visibly reduced at all three concentrations (**Fig. 3a-e**). In each case some fully EGFP positive embryos were recovered, an effect that may be due to a variable position of eggs within the cuvette. For this reason, we suspect that careful mixing of eggs and shRNA within the cuvette can improve efficiency. To quantify the results at a molecular level, we analyzed relative EGFP mRNA levels by qPCR. In keeping with the image analysis, we observed a marked reduction in target mRNA levels at all three shRNA concentrations (**Fig. 3f**).

**Figure 3.**
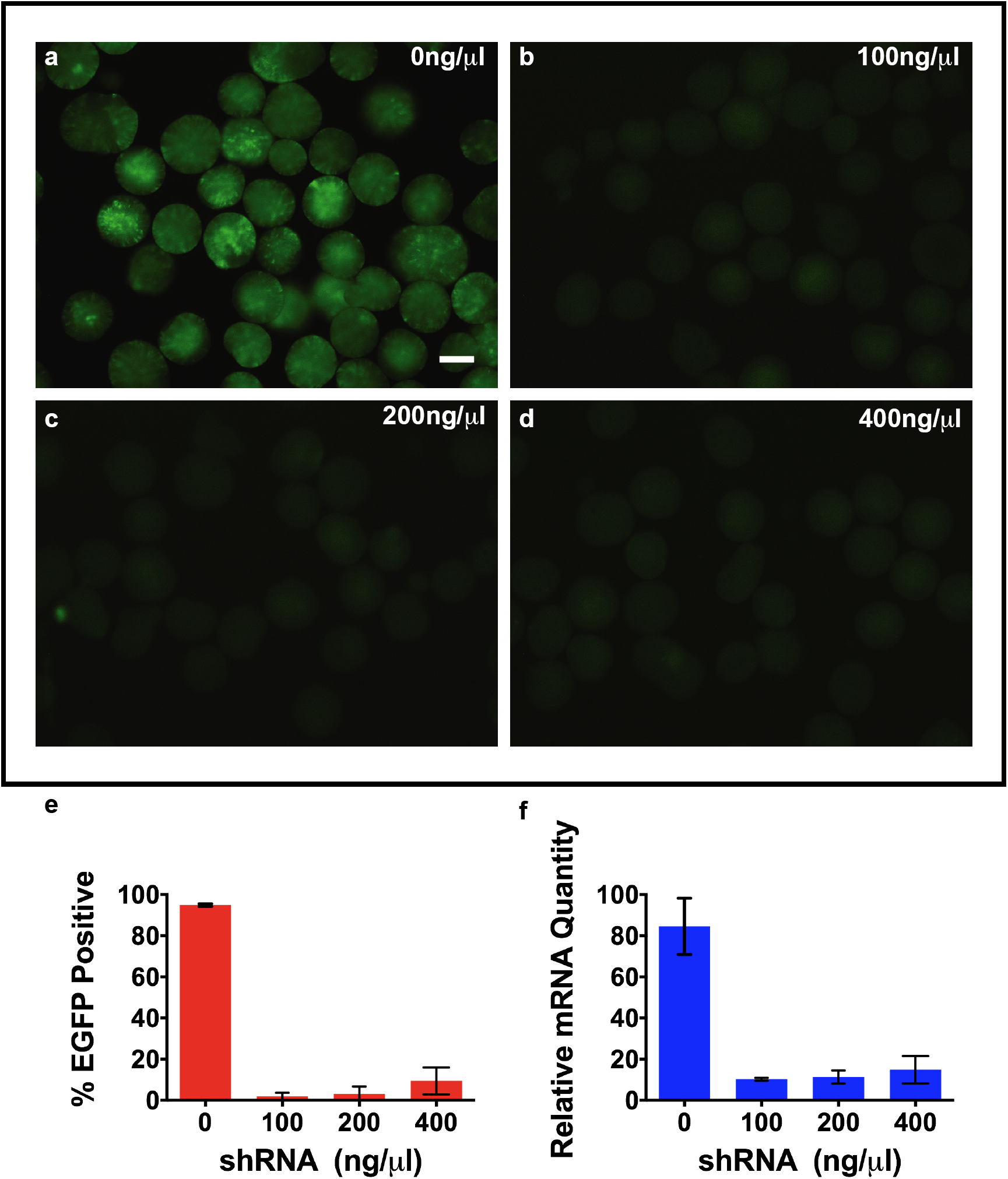
Quantification of the of gene expression levels and number of embryos that are positive for each dose after electroporation into *Actin-EGFP* transgenic animals. (**a**) 0ng/μl, (**b**)100ng/μl (**c**) 200ng/μl (**d**) 400ng/μl indicate the doses of EGFP shRNA. Images were acquired 48 hours after electroporation. (**e**) Gene expression levels and the number of embryos that correspond to EGFP signal after electroporation. (Scale bar = 250 μm)

In addition to knockdown of an exogenous GFP reporter gene, we also used electroporation to deliver shRNAs targeting four endogenous loci (*Nvβ–catenin*, *NvChordin*, *NvBMP5/8* and *NvAnthox1a*). Electroporation of shRNA targeting *Nvβ-catenin* blocked gastrulation and disrupted cell-cell cohesion by approximately 20 hours post-fertilization (hpf) with 99% phenotypic penetrance (**Fig. 4**). This finding is similar but more severe than the published morpholino phenotypes^26^. In addition, electroporation of shRNAs against *NvChordin* and *NvBMP5/8* reproduced expected loss of function phenotypes at 74% and 52% penetrance, respectively^27^. In both cases, knockdown resulted in animals with highly elongated body columns and reduced tentacles (**Fig. 5**). Also in keeping with previous data, electroporation of shRNA targeting *NvAnthox1a* resulted in the expected tentacle bifurcation phenotype with 64% penetrance **(Fig. 6;** He *et al*., *in press*). Combined, these results indicate that electroporation-mediated shRNA knockdown is a specific and potent tool for rapid analysis of gene function in *Nematostella*.

**Figure 4.**
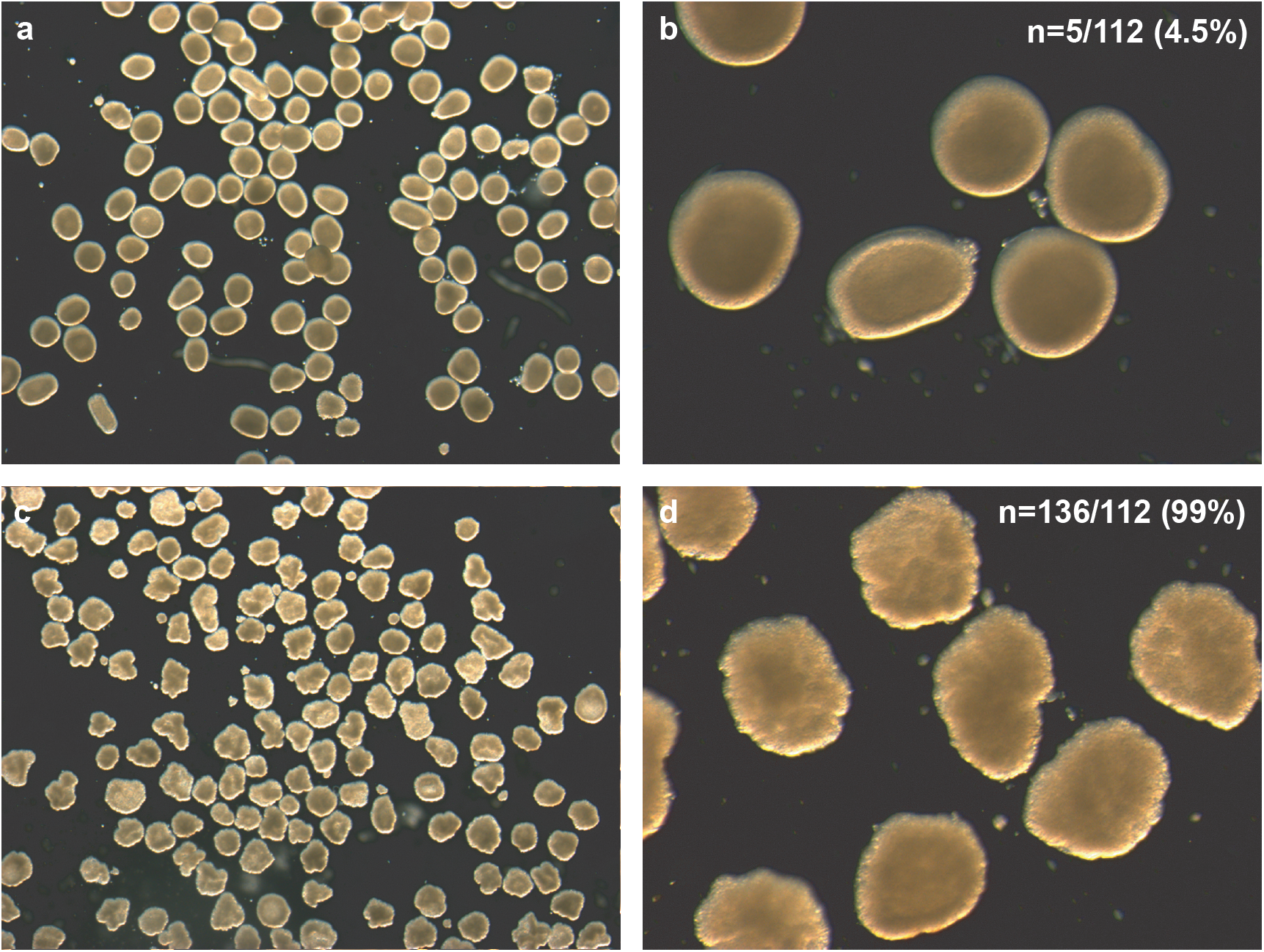
Phenotypes of (**a**) control shRNA (300ng/μl) and (**b**) shRNA targeting *Nv*β*-catenin* gene product (300ng/μl) after electroporation. Close up images of (**c**) control shRNA (**d**) *Nv*β**-***catenin* shRNA. Images were acquired 72 hours after electroporation.

**Figure 5.**
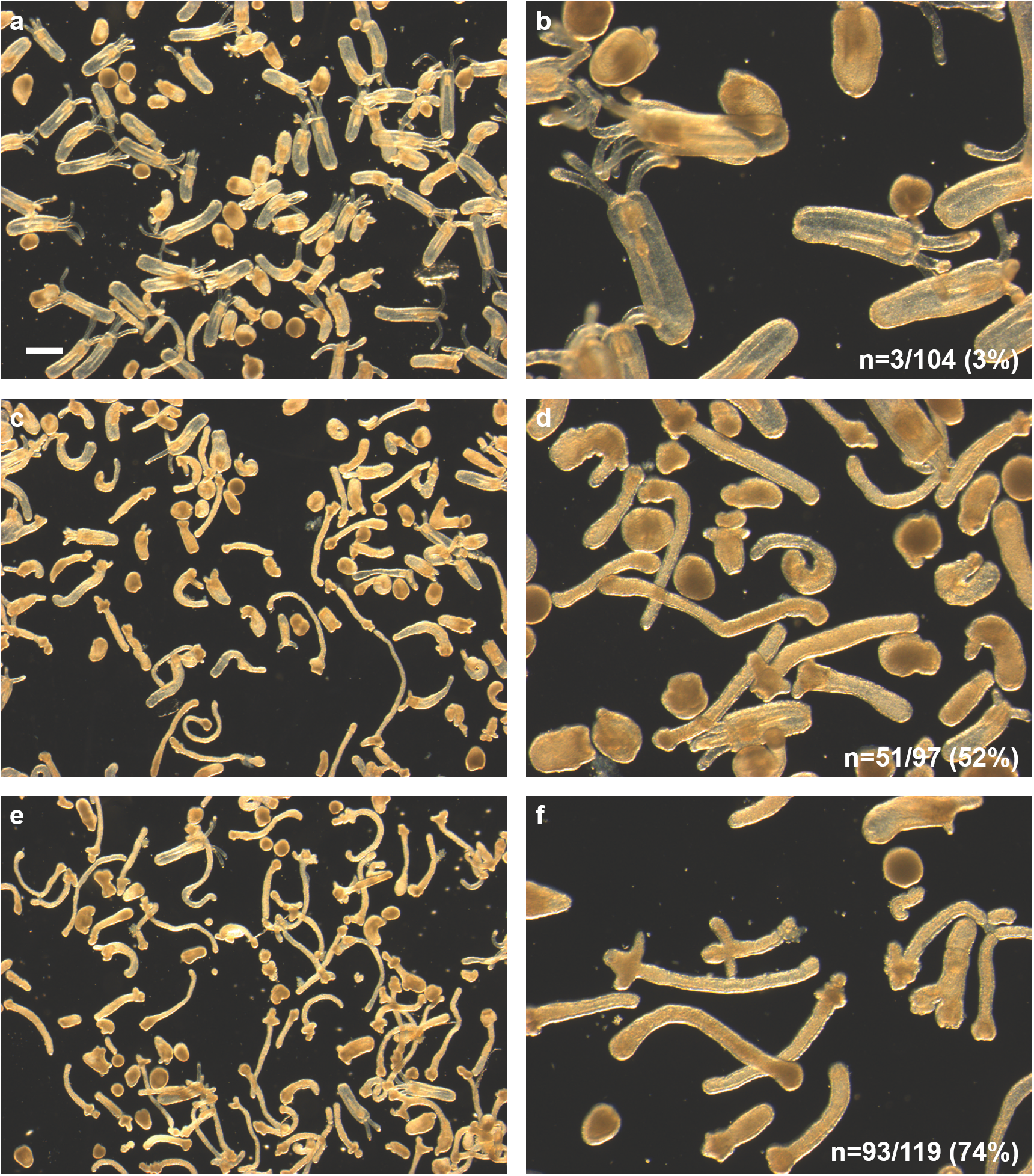
Phenotypes of (**a**) Scramble shRNA (300ng/μl) and (**b**) Scramble shRNA high magnification. (**c**) *NvBMP5/8* shRNA (300ng/μl) and (**d**) *NvBMP5/8* shRNA high magnification (**e**) *Nvchordin* shRNA (300ng/μl) (**f**) *Nvchordin* shRNA high magnification. Numbers indicate counts and percentage of knockdown versus wildtype phenotypes in images (**a**), (**b**) and (**c**) respectively. Images were acquired 6 days after electroporation. (Scale Bar = 250 μm)

**Figure 6.**
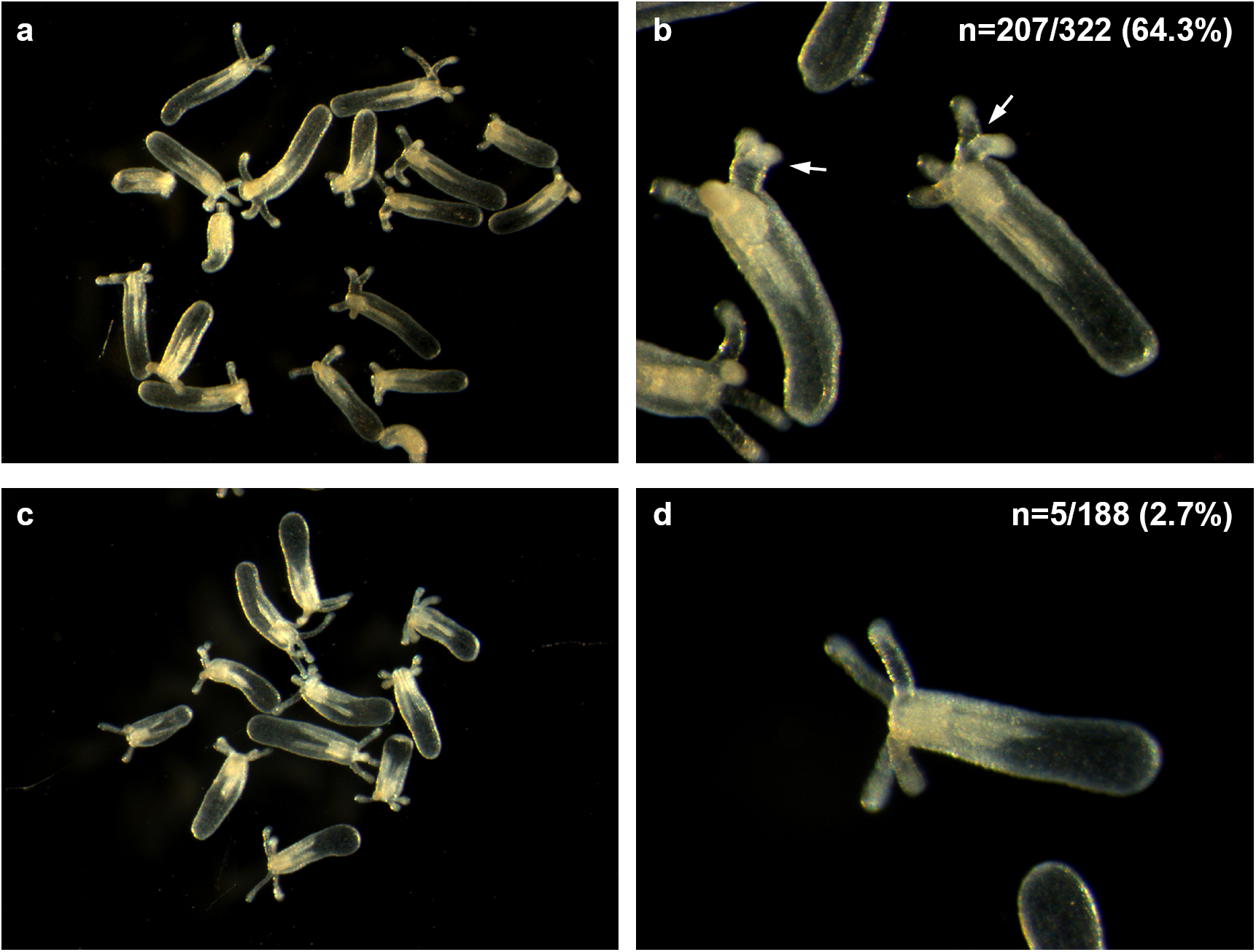
Phenotypes of (**a**) *NvAnthox1a* shRNA (900ng/μl) and (**b**) *NvAnthox1a* shRNA with high magnification. (**c**) Scramble shRNA (900ng/μl) and (**d**) Scramble shRNA with high magnification. Images were acquired 9 days after electroporation.

While high doses of shRNA are tolerated, knockdown-dependent lethal phenotypes are frequently observed at much lower doses. To determine the phenotype for an unknown gene, we recommend a dose response, starting with a high concentration i.e. 500ng/μl and a lower concentration, i.e. 200 ng/μl. Above, we tested up to 900 ng/μl shRNA without any observed toxicity **(Fig. 6)**, suggesting that a broad range of shRNA concentrations can be used knock down a target of interest. We further recommend that a scrambled shRNA as well as shRNAs that produce known phenotypes be used as negative and positive controls. As presented here, *β-catenin*, *BMP5/8* and *Chordin* shRNAs all produce obvious phenotypes and can be used to validate both electroporation and knockdown efficiency. Additionally, non-electroporated controls should be included to evaluate general fertilization efficiency and viability for each experiment. Importantly, the distinct phenotypes of *NvChordin*, *NvBMP5/8*, *Nvβ-catenin* can be used as positive controls for phenotypic analysis at different time points. For example, knockdown of Nv*β-catenin* can be used as a positive control for early-expressed genes at 24hpf, whereas *NvBMP5/8*, *NvChordin* and *NvAnthox1a* can be used as a positive control for later phenotypes that manifest in the range of 5-7 days post-fertilization.

## CONCLUSION

Electroporation is an efficient and scalable method to deliver shRNA into *Nematostella* eggs. This procedure is reproducible and should provide a similar knockdown efficiency as microinjection in a significantly reduced timeframe. We recommend 500ng/μl and 200ng/μl as starting shRNA concentrations. In our experience, shRNA should efficiently knock down a gene of interest at 500ng/μl.

## PROTOCOL

### Reagents

#### For shRNA synthesis and purification

- Forward and reverse primers for template annealing.
- AmpliScribe™ T7-*Flash*™ transcription kit (Lucigen, Inc).
- Direct-zol™ Miniprep Plus (Zymo Research, R2070).
- Tri-Reagent (Ambion, Inc).
- Nuclease-free water.
- Oligonucleotides (Integrated DNA Technologies).

### Equipment

#### For shRNA synthesis and purification

- Thermal Cycler (Bio-rad DNA engine) or equivalent.
- Benchtop Centrifuge (Eppendorf, 5430R) or equivalent.
- Microcentrifuge tubes (Denville, Cat. C2170).

#### For egg preparation

- Nutator (Clay Adams).
- Conical bottom tubes (15ml; Grenier Bio-One, Cat. 188261).
- Disposable Graduated Transfer Pipettes, (VWR, Cat. No. 414004-014).
- Crystallizing Dishes, (Kimax Kimble, Cat. No. 23000).
- L-Cysteine (Sigma, 168149).

#### For electroporation

- ECM 830 Electro Squire Porator (BTX™, Cat. No 450052).
- Electroporation Cuvettes (4mm; Mirus, Cat. No. MIR50123).
- Gene Pulser Xcell ShockPod Cuvette Chamber (Bio-rad, Cat. No. 1652669).
- Laboratory wash bottle (500ml; VWR 47750-710).
- Conical tubes (15ml; Grenier Bio-One, cat. no. 188261).
- Microcentrifuge tubes (Denville, Cat. No.C2110).
- Plastic petri dishes (60 mm X 15 mm) (Falcon,351007).
- Disposable Graduated Transfer Pipets (VWR North America, Cat. No. 414004-014).
- Microcentrifuge transfer pipet (RPI; Cat. No. 147500).
- Single edge razor (VWR, Cat. No. 55411-050).

### For image acquisition

#### Inverted fluorescence microscope

Images were acquired on a Zeiss Axiovert 200M with Fluar 5X 0.25NA equipped with Micro-Manager1. Processing was done in Fiji. Individual embryos were segmented using custom thresholding, separated using watershed and mean intensity quantified. Processing macros and plugins can be found at https://github.com/jouyun/^28^. Brightfield microscope images were acquired in a Leica MZ16 Stereomicroscope. Embryos were counted and distinguished by using thresholding, segmentation using watershed and particle analysis in Fiji.

#### For RT-qPCR

We used the QuantStudio(TM) 7 Flex System (Thermo Fisher). Promega ImProm-II™ Reverse Transcriptase (cat. no. A3801) was used for RT reactions. https://www.promega.com/products/pcr/rt-pcr/improm_ii-reverse-transcriptase/?catNum=A3801 qPCR was performed with NEB Luna® Universal qPCR Master Mix (M3003) at10µl reaction volume. The cycles were according to the supplier’s manual. https://www.neb.com/products/m3003-luna-universal-qpcr-master-mix#Product%20Information

The following Primers were used for the qPCR reaction:

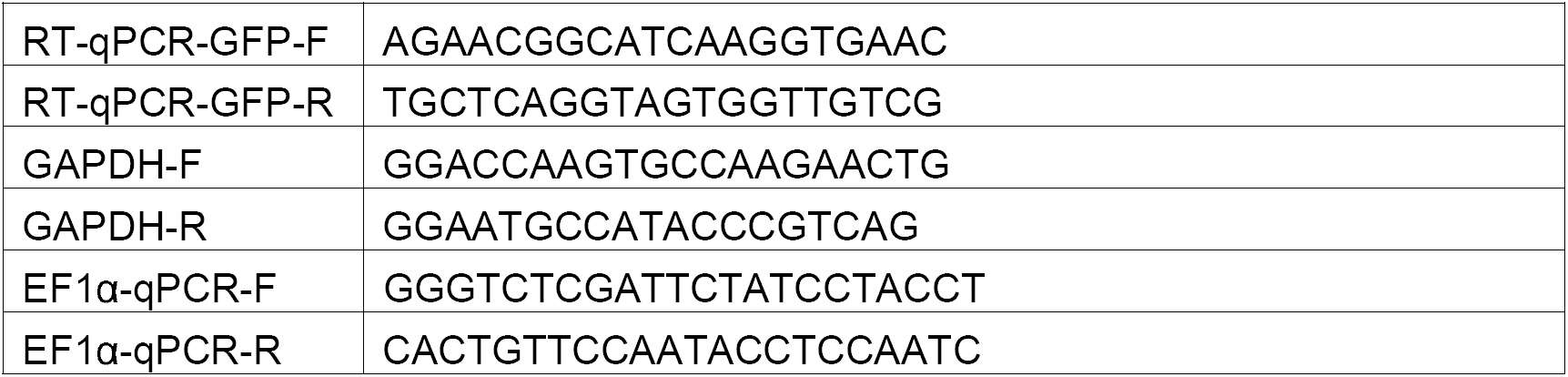

### Reagent preparation

#### *Nematostella* artificial saltwater (ASW), 12 ppt

Dissolve 1.2g of artificial sea salt (Instant Ocean) into deionized water to a final volume of 1L. ASW can be stored at room temperature for approximately one month.

#### Egg dejellying solution

Dissolve 1g of L-Cysteine hydrochloride into 25ml ASW. Add 5 drops of 5M NaOH to adjust the pH and place on a rocker until L-Cysteine is dissolved. The solution must be freshly prepared and used immediately before de-jellying.

#### Egg suspension medium (15% (wt/vol) Ficoll PM400)

In a 50ml Falcon tube, dissolve 3g Ficoll PM400 in 15ml ASW. Vortex for 2 minutes and shake in a rotary shaker for 30 minutes. Adjust the volume to 20ml by adding ASW. Vortex for 2 minutes, let the tube settle. Solution will initially look cloudy but will gradually clear within ten minutes.

#### Note

Ficoll dissolves in ASW slowly at room temperature. When dissolved, the solution should be clear and slightly viscous. If Ficoll doesn’t dissolve well, incubate the Ficoll in ASW for 20 minutes at 50°C and vortex every 5 minutes during incubation.

### Equipment setup

#### Electroporation

**1.** Connect the Gene Pulser Xcell ShockPod Cuvette Chamber (Bio-rad) cable.

**2.** Turn on the ECM830 electroporator. Wait for the device to boot and push the knob to select the parameters (i.e. volts, pulse duration, number of pulses and pulse interval). Adjust settings as follows: 50 volts, 25 milliseconds, 1 pulse and 500 milliseconds pulse interval. The device should be set ready for electroporation. Most comparable laboratory electroporators should be sufficient to achieve these conditions in a 4mm gap cuvette.

#### Cleaning and re-use of Electroporation Cuvettes

• 4mm cuvettes can be reused at least 10 times if washed and dried thoroughly after each electroporation. Clean the cuvettes with de-ionized water followed by a second wash with 100% ethanol. After the ethanol washes, dry at room temperature or 37°C. Avoid using detergents to wash the cuvette. Cuvettes can be kept at room temperature indefinitely.

## DETAILED PROCEDURE

### (A) shRNA design TIMING (1-3 hours)

**1.** Retrieve cDNA sequences of the target genes from Joint Genome Institute’s *Nematostella* genome portal. (https://genome.jgi.doe.gov/pages/blast-query.jsf?db=Nemve1)

**2.** Open shWizard software. http://www.invivogen.com/sirnawizard/index.php

**3.** Set motif size to 19 nucleotides and paste cDNA sequence of interest.

**4.** Click search button, several candidate target sequences will be generated.

**5.** Select two or three motifs.

**6.** Click design button and retrieve the sense and anti-sense sequences. **See Table** Note: Motifs must have reasonable complexity, a CG content between 40-55% and must have a relatively low CG content at the 3’ end. Total length of the template is 66 bases.

**7.** BLAST candidate sequences to JGI website for the *Nematostella* Genome database. Eliminate sequences with multiple hits.

**8.** In the program settings box, set the “expect” box to 1.02E-2 for short sequences.

**9.** Copy sequences with a single match in the *Nematostella* genome.

**10.** Design the IVT template by inserting the motifs according to the highlighted positions indicated in **Table 1, Row 1.**

Note: It is essential that the final DNA template is 66 base pairs long with the T7 promoter at the 5’ end, which is necessary for IVT. The final DNA template length must be 66 base pairs to avoid misfolding of the synthesized shRNA. Use mfold software from the University of Albany (http://unafold.rna.albany.edu/?q=mfold) to analyze the secondary structure of the shRNA to ensure that there is only one conformer. ^29^

### (B) shRNA synthesis and production TIMING (8 hours)

**11.** To construct the IVT template, first order forward and reverse primers as described in Step 10.

Note: The final template should have 19 base sense and anti-sense targeting sequences as described in (**Table 1, Row 1).**

**12.** Dissolve the primers in nuclease free water to a final concentration of 50μM.

**13.** In a PCR reaction tube, mix 5μl of forward and reverse primers to a final volume of 10μl.

Note: Mix with caution. Avoid mixing slightly higher or lower volumes of each primer as this changes the final duplex concentration for the IVT reaction.

**14.** In a thermal cycler, denature the primer mix at 98°C for 5 minutes and immediately cool down to 24°C to anneal the forward and reverse primers.

**15.** Use Ampliscribe™ T7-Flash™ Transcription Kit for the following steps.

**16.** Prepare the IVT reaction mix at room temperature. For a 20μl reaction:

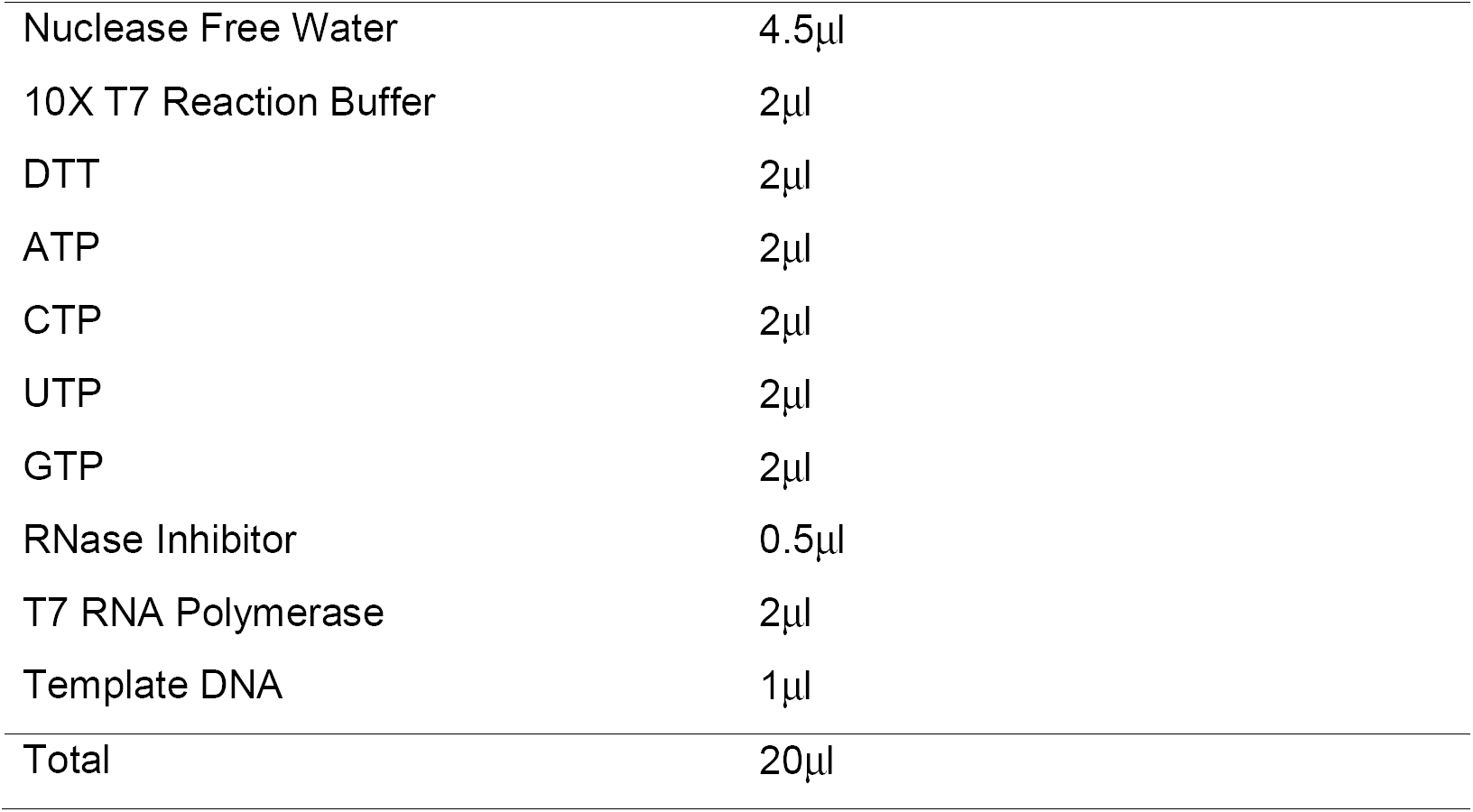

Note: Kit components may precipitate after preparation of the reaction mix. If this occurs, heat the components at 37°C for 10 minutes.

**17.** Allow the IVT reaction to continue for 7 hours at 37°C. The reaction can be continued for 15 hours or overnight if desired. However, no significant increase in yield will be achieved.

**18.** At this point, the reaction mixture can be temporarily stored at 4°C as needed. Move to −20°C for storage longer than 24 hours.

**19.** To eliminate the IVT DNA template, add 1μl of RNase-free DNase to the reaction mix and incubate for 15 minutes at 37°C.

**20.** At room temperature, dilute the reaction product 1:10 with DEPC-treated water. Add an equal volume of ethanol (95-100%) as described below.

|  |  |
| --- | --- |
| T7 Reaction Mix | 20µl |
| DEPC treated water | 180µl |
| 100% Ethanol | 200µl |
| Total | 400µl |

**21.** Purify shRNA from the mixture using a Zymogen RNA Plus Miniprep spin column according to the manufacturer’s instructions.

**22.** Following elution in nuclease-free water, measure shRNA concentration by Nanodrop or an equivalent spectrophotometer. The OD (260/280) ratio should be above 1.80.

Note: The OD (260/280) reading can be above the measurable range. Dilute 1μl of eluted product with 99μl nuclease-free water to obtain accurate readings.

**23.** As an optional control, run 1μl of the eluent in an agarose gel to confirm the spectrophotometer results. The IVT Reaction Mix can be stored up to 2 months at −20°C.

### (C) Dejellying and Preparation of Eggs for electroporation TIMING (40 minutes)

**24.** Induce spawning of adult males and females in separate dishes according to the previously-described protocol^15^.

**25.** Collect unfertilized egg sacs in 15ml conical tubes and distribute 2ml of eggs per tube. Add 12ml L-Cysteine solution (to the 14ml mark) using a 1ml pipette.

**26.** Mix tubes on a nutator for 10 minutes.

**27.** Let eggs settle to the bottom for approximately 1 minute. Remove water down to 2 milliliters and replace with fresh ASW.

**28.** Gently invert the tube to suspend eggs. Perform 3 washes with ASW.

**Note:** Transfer pipettes can damage the eggs. To avoid this, cut off the end of the pipet to increase the diameter of the opening.

**29.** After the final wash, remove water and use a pipette to resuspend eggs in ASW. Transfer eggs into a 90×50 crystallizing dish containing ASW (12 p.p.t.). Keep at 18°C until electroporation.

Note: Keeping the sperm at 18°C should lengthen stability, but use within 2 hours is recommended. Eggs can be kept for up to 4 hours at 18°C after dejellying. Sperm can be kept for 2 hours at 23°C.

### (D) Electroporation procedure TIMING (20 minutes)

**30.** Use a transfer pipette to move eggs from the crystalizing dish to a 15ml conical tube. Allow eggs to settle and then transfer concentrated eggs to a fresh tube with a volume for the desired number of electroporations.

Note: Each electroporation requires 100μl of eggs suspended in 15% Ficoll ASW. Hence for 10 electroporations, transfer concentrated eggs to a 1.5 ml microcentrifuge tube. For larger-scale experiments, transfer to a 15ml conical tube.

**31.** Remove ASW and replace with ASW containing 15% Ficoll PM 4000.

**32.** Suspend the eggs by pipetting the mixture with a Gelatin- or Ficoll-coated 1ml transfer pipet.

Note: Eggs can settle over time. (i.e. 5 minutes). If this happens gently tap the tube to re-suspend the eggs. Prepare and use coated plastic polypropylene pipettes for all transfers. Eggs stick to pipette tips if the tips are not coated with Ficoll or Gelatin.

**33.** Use a razor to cut a 200μl pipette tip to increase the diameter.

**34.** Carefully transfer suspended eggs into a 4mm cuvette using a 200μl pipette.

Note: Avoid touching the walls of the cuvette with the tip. Eggs that are stuck to the sides of the chamber or otherwise excluded from the main reaction volume will not be subjected to electroporation.

**35.** Add purified shRNA directly to the main reaction volume in the electroporation cuvette.

**36.** Gently mix shRNA and eggs by shaking side to side.

Note: Shake 10 to 30 seconds to ensure complete mixing before electroporation.

**37.** Turn on ECM 830 Electroporator or equivalent device.

**38.** The device should be in low voltage mode (LV) as displayed on the LCD screen. Set the electroporation conditions to 50 volts, 25 msec pulse duration, 1 pulse and 500 msec pulse interval.

**39.** Insert the electroporation cuvette into the Bio-Rad Shock Pod cuvette receiver.

**40.** Press start button to electroporate. A short buzzing sound will be heard. The LCD screen will display the actual voltage, pulse duration and number of pulses. i.e. 46 volts 25msec, 1 pulse.

Note: Each experiment should include controls with and without electroporation. These controls will provide information about the general fertilization rate for that day and embryo survival after electroporation along with the shRNA controls.

**41.** Remove the cuvette. On close examination, small bubbles should be observed on the walls of the cuvette.

**42.** Immediately use a transfer pipette to fill the cuvette with *Nematostella* sperm water from a dish of sex-sorted males.

**43.** Add enough sperm water to cover the bottom of a plastic petri dish (60 x 15 mm).

**44.** Transfer the entire volume of electroporated eggs into this petri dish using the same transfer pipette.

Note: Make sure the developing embryos are well dispersed in the (60 x 15mm) petri dish for incubation. Embryos can stick to each other and may form conjoined animals.

**45.** Incubate the plates at 18°C or 23°C overnight.

**46.** After 24 hours remove debris and perform 2 or 3 exchanges of ASW. Using a transfer pipette, gently transfer the embryos to a new clean plate filled with ASW.

**47.** Monitor development relative to controls and perform more detailed phenotypic studies as the desired stage.

## ACKNOWLEDGEMENTS

We thank Aissam Ikmi for sharing the *NvActin*-EGFP transgenic line, and Lacey Ellington for reading and editing the manuscript. We also thank Florencia Del Viso, Katerina Ragkousi, Eric Hill and Gibson Lab members for their recommendations toward the development of this protocol. We also thank the Stowers Institute for Medical Research Reptile and Aquatics facility for animal care and Hang Yin and Abby Dreyer for administrative assistance.

## AUTHOR CONTRIBUTIONS

M.G. conceived the paper. A.K. developed the electroporation protocol. S.H. developed the method for shRNA-based gene knockdown. A.K. and S.H performed the experiments. A.K. S.H. C.C. and S.M. generated the data.

**Table 2:** Possible problems and solutions.

| **Step** | **Problem** | **Reason** | **Solution** |
| --- | --- | --- | --- |
| 16 | Precipitate observed during IVT reaction assembly. | The IVT reagents were not warm enough. | Incubate the reaction mix before enzyme addition at 37°C for 10 minutes. If the solution is clear add T7 RNA Polymerase. |
| 22 | Very low shRNA yield. | IVT mix precipitated after enzyme addition. | Inspect the IVT reaction mix for precipitation. If precipitation is observed, after enzyme addition, heat the IVT mix at 37°C for 10 minutes. If there is precipitation, discard the tube and make a new reaction mix. |
| 32 | Rapid Resettlement of Eggs suspended in ASW 15% Ficoll | Excess water is left in the tube before ASW - 15% Ficoll addition. | Resuspend by gentle tapping before transfer to the cuvette |
| 42 | Very Low Fertilization Rate | Reduced sperm quality. Poor Contact of Sperm with Egg. | Add fresh sperm when possible. Keep sperm at 18°C until use. Gently swirl the plate to mix sperm with electroporated eggs |
| 44 | Embryos and primary polyps are fused and look unhealthy. | Embryos clumped at the center of the plate. | Disperse eggs carefully and avoid crowding at the center of the petri dish after electroporation. Move slowly while carrying the plates to a different location. |
| 47 | Very low numbers of embryos with known phenotype. | shRNA is not mixed well. Low quality shRNA. shRNA dose is low. | Check the quality of shRNA by running the IVT product in an agarose gel. Thoroughly mix shRNA and egg suspension by gentle shaking the cuvette for 30 seconds. If phenotype is present, electroporate a higher concentration of shRNA. |
| 47 | Unexpectedly high number of normal embryos in positive control dishes. | Pipetting error. Eggs stick to the non-conductive regions of the cuvette during embryo and shRNA transfer.<br>If Sperm source is a mixed-sex dish, it might be contaminated with eggs | Wipe the cuvettes with Kim Wipes before transferring eggs to the cuvette.<br>If the sperm is originated from a mixed dish, inspect sperm dish for egg contamination before use. |

## REFERENCES

1 Layden, M. J., Rentzsch, F. & Rottinger, E. The rise of the starlet sea anemone Nematostella vectensis as a model system to investigate development and regeneration. Wiley Interdiscip Rev Dev Biol 5, 408–428, doi:10.1002/wdev.222 (2016).

2 Christiaen, L., Wagner, E., Shi, W. & Levine, M. Isolation of individual cells and tissues from electroporated sea squirt (Ciona) embryos by fluorescence-activated cell sorting (FACS). Cold Spring Harb Protoc 2009, pdb prot5349, doi:10.1101/pdb.prot5349 (2009).

3 Genikhovich, G. & Technau, U. The starlet sea anemone Nematostella vectensis: an anthozoan model organism for studies in comparative genomics and functional evolutionary developmental biology. Cold Spring Harb Protoc 2009, pdb emo129, doi:10.1101/pdb.emo129 (2009).

4 Putnam, N. H. et al. Sea anemone genome reveals ancestral eumetazoan gene repertoire and genomic organization. Science 317, 86–94, doi:10.1126/science.1139158 (2007).

5 Guder, C. et al. The Wnt code: cnidarians signal the way. Oncogene 25, 7450–7460, doi:10.1038/sj.onc.1210052 (2006).

6 Marlow, H., Roettinger, E., Boekhout, M. & Martindale, M. Q. Functional roles of Notch signaling in the cnidarian Nematostella vectensis. Dev Biol 362, 295–308, doi:10.1016/j.ydbio.2011.11.012 (2012).

7 Fritz, A. E., Ikmi, A., Seidel, C., Paulson, A. & Gibson, M. C. Mechanisms of tentacle morphogenesis in the sea anemone Nematostella vectensis. Development 140, 2212–2223, doi:10.1242/dev.088260 (2013).

8 Saina, M., Genikhovich, G., Renfer, E. & Technau, U. BMPs and chordin regulate patterning of the directive axis in a sea anemone. Proc Natl Acad Sci U S A 106, 18592–18597, doi:10.1073/pnas.0900151106 (2009).

9 Matus, D. Q., Thomsen, G. H. & Martindale, M. Q. FGF signaling in gastrulation and neural development in Nematostella vectensis, an anthozoan cnidarian. Dev Genes Evol 217, 137–148, doi:10.1007/s00427-006-0122-3 (2007).

10 Rentzsch, F., Fritzenwanker, J. H., Scholz, C. B. & Technau, U. FGF signalling controls formation of the apical sensory organ in the cnidarian Nematostella vectensis. Development 135, 1761–1769, doi:10.1242/dev.020784 (2008).

11 Matus, D. Q., Magie, C. R., Pang, K., Martindale, M. Q. & Thomsen, G. H. The Hedgehog gene family of the cnidarian, Nematostella vectensis, and implications for understanding metazoan Hedgehog pathway evolution. Dev Biol 313, 501–518, doi:10.1016/j.ydbio.2007.09.032 (2008).

12 Sullivan, J. C., Reitzel, A. M. & Finnerty, J. R. A high percentage of introns in human genes were present early in animal evolution: evidence from the basal metazoan Nematostella vectensis. Genome Inform 17, 219–229 (2006).

13 Zimek, A. & Weber, K. In contrast to the nematode and fruit fly all 9 intron positions of the sea anemone lamin gene are conserved in human lamin genes. Eur J Cell Biol 87, 305–309, doi:10.1016/j.ejcb.2008.01.003 (2008).

14 Hand, C. & Uhlinger, K. R. The Culture, Sexual and Asexual Reproduction, and Growth of the Sea Anemone Nematostella vectensis. Biol Bull 182, 169–176, doi:10.2307/1542110 (1992).

15 Genikhovich, G. & Technau, U. Induction of spawning in the starlet sea anemone Nematostella vectensis, in vitro fertilization of gametes, and dejellying of zygotes. Cold Spring Harb Protoc 2009, pdb prot5281, doi:10.1101/pdb.prot5281 (2009).

16 Stefanik, D. J., Wolenski, F. S., Friedman, L. E., Gilmore, T. D. & Finnerty, J. R. Isolation of DNA, RNA and protein from the starlet sea anemone Nematostella vectensis. Nat Protoc 8, 892–899, doi:10.1038/nprot.2012.151 (2013).

17 Wolenski, F. S., Layden, M. J., Martindale, M. Q., Gilmore, T. D. & Finnerty, J. R. Characterizing the spatiotemporal expression of RNAs and proteins in the starlet sea anemone, Nematostella vectensis. Nat Protoc 8, 900–915, doi:10.1038/nprot.2013.014 (2013).

18 Genikhovich, G. & Technau, U. In situ hybridization of starlet sea anemone (Nematostella vectensis) embryos, larvae, and polyps. Cold Spring Harb Protoc 2009, |ppdb prot5282, doi:10.1101/pdb.prot5282 (2009).

19 Layden, M. J., Rottinger, E., Wolenski, F. S., Gilmore, T. D. & Martindale, M. Q. Microinjection of mRNA or morpholinos for reverse genetic analysis in the starlet sea anemone, Nematostella vectensis. Nat Protoc 8, 924–934, doi:10.1038/nprot.2013.009 (2013).

20 Renfer, E. & Technau, U. Meganuclease-assisted generation of stable transgenics in the sea anemone Nematostella vectensis. Nat Protoc 12, 1844–1854, doi:10.1038/nprot.2017.075 (2017).

21 Ikmi, A., McKinney, S. A., Delventhal, K. M. & Gibson, M. C. TALEN and CRISPR/Cas9-mediated genome editing in the early-branching metazoan Nematostella vectensis. Nat Commun 5, 5486, doi:10.1038/ncomms6486 (2014).

22 Brennecke, T., Gellner, K. & Bosch, T. C. The lack of a stress response in Hydra oligactis is due to reduced hsp70 mRNA stability. Eur J Biochem 255, 703–709 (1998).

23 Kari, W., Zeng, F., Zitzelsberger, L., Will, J. & Rothbacher, U. Embryo Microinjection and Electroporation in the Chordate Ciona intestinalis. J Vis Exp, doi:10.3791/54313 (2016).

24 Vierra, D. A. & Irvine, S. Q. Optimized conditions for transgenesis of the ascidian Ciona using square wave electroporation. Dev Genes Evol 222, 55–61, doi:10.1007/s00427-011-0386-0 (2012).

25 Ge, Q. et al. Minimal-length short hairpin RNAs: the relationship of structure and RNAi activity. RNA 16, 106–117, doi:10.1261/rna.1894510 (2010).

26 Watanabe, H. et al. Sequential actions of beta-catenin and Bmp pattern the oral nerve net in Nematostella vectensis. Nat Commun 5, 5536, doi:10.1038/ncomms6536 (2014).

27 Leclere, L. & Rentzsch, F. RGM regulates BMP-mediated secondary axis formation in the sea anemone Nematostella vectensis. Cell Rep 9, 1921–1930, doi:10.1016/j.celrep.2014.11.009 (2014).

28 Edelstein, A. D. et al. Advanced methods of microscope control using muManager software. J Biol Methods 1, doi:10.14440/jbm.2014.36 (2014).

29 Rouillard, J. M., Zuker, M. & Gulari, E. OligoArray 2.0: design of oligonucleotide probes for DNA microarrays using a thermodynamic approach. Nucleic Acids Res 31, 3057–3062 (2003).

